# An Image Processing Tool for Automated Quantification of Bacterial Burdens in Zebrafish Larvae

**DOI:** 10.1101/2024.08.16.608298

**Authors:** Naoya Yamaguchi, Hideo Otsuna, Michal Eisenberg-Bord, Lalita Ramakrishnan

## Abstract

Zebrafish larvae are used to model the pathogenesis of multiple bacteria. This transparent model offers the unique advantage of allowing quantification of fluorescent bacterial burdens (fluorescent pixel counts: FPC) in vivo by facile microscopical methods, replacing enumeration of bacteria using time-intensive plating of lysates on bacteriological media. Accurate FPC measurements require laborious manual image processing to mark the outside borders of the animals so as to delineate the bacteria inside the animals from those in the culture medium that they are in. Here, we have developed an automated ImageJ/Fiji-based macro that accurately detect the outside borders of *Mycobacterium marinum*-infected larvae.

## Introduction

The zebrafish larva has come into its own as a model for bacterial pathogenesis and drug discovery (1-4). In addition to its genetic amenability, its optical transparency allows real-time visualization of infection. Husbandry is facile; larvae can be maintained for two weeks in petri dishes without feeding or media changes, allowing the addition of drugs to be tested to the media.

Determining in vivo bacterial burdens is a core component of infectious diseases research. The classical method — enumerating bacteria from lysed animals or tissues following plating on bacteriological media — is laborious and time-consuming. Visible colonies may take days to weeks to appear, depending on the pathogen. Furthermore, serial assessments cannot be made in the same animals.

We overcame this problem by developing a microscopical method to determine relative burdens of fluorescent bacteria by enumerating fluorescence pixel counts (FPC) within the outlines of larval bodies (5-7). 96-well plates containing the larvae are imaged using an inverted fluorescence microscope with a motorized stage, allowing hundreds of measurements to be completed within an hour. Serial imaging of the larvae allows monitoring of infection progression and testing drug efficacy. It can be used to study nonculturable pathogens such as *Mycobacterium leprae* rendered fluorescent using dyes (8).

However, the throughput of the FPC method is hindered by the need for a manual image analysis step: fluorescent bacteria shed from skin and other debris create bright spots outside the body, which can interfere with the thresholding process during image analysis (**Supplemental Material 1**, Supplemental Figure 1). Precise analysis of bacteria within the animal requires time-consuming, labor-intensive manual blacking out of the space outside of the larva. Furthermore, the manual method requires a threshold value determined by each user, making for inter-user variability in the analysis.

Here, we have developed an Image J/Fiji-based macro to automatically identify the larval body contour and bacterial spots within it and validated it using zebrafish infected with fluorescent *Mycobacterium marinum* (Mm).

## Results and Discussion

We developed a Graphical User Interface (GUI)-based Image J/Fiji macro (9) (Fig. 1A). This macro is designed to allow users to collect various features of bacterial foci from the dataset (Fig. 1B; **Supplemental Material 2)**. An input image is taken using a conventional widefield fluorescence microscope utilizing both fluorescence and dim transmitted light to simultaneously delineate the larval body and the fluorescent bacteria within it (Fig. 1C, left). The macro initially determines the outer border of the zebrafish larva from the 96-well image (Fig. 1C, middle). While this process is not perfect, it is sufficient for most of the images that we tested to proceed to the next step. Once the larval “object” is determined, fluorescence bacterial foci within the larval object are detected, and output parameters are measured (Fig. 1C, right). The individual image processing steps are summarized in **Supplemental Material 3**. Additionally, the macro allows users to explore the “Segmentation sensitivity” to identify the optimal segmentation for their images. Typically, running the macro with a lower Segmentation sensitivity results in segmenting the outer border of the larva closer to the larval body (Fig.1D; **Supplemental Material 3**).

**Figure 1:**
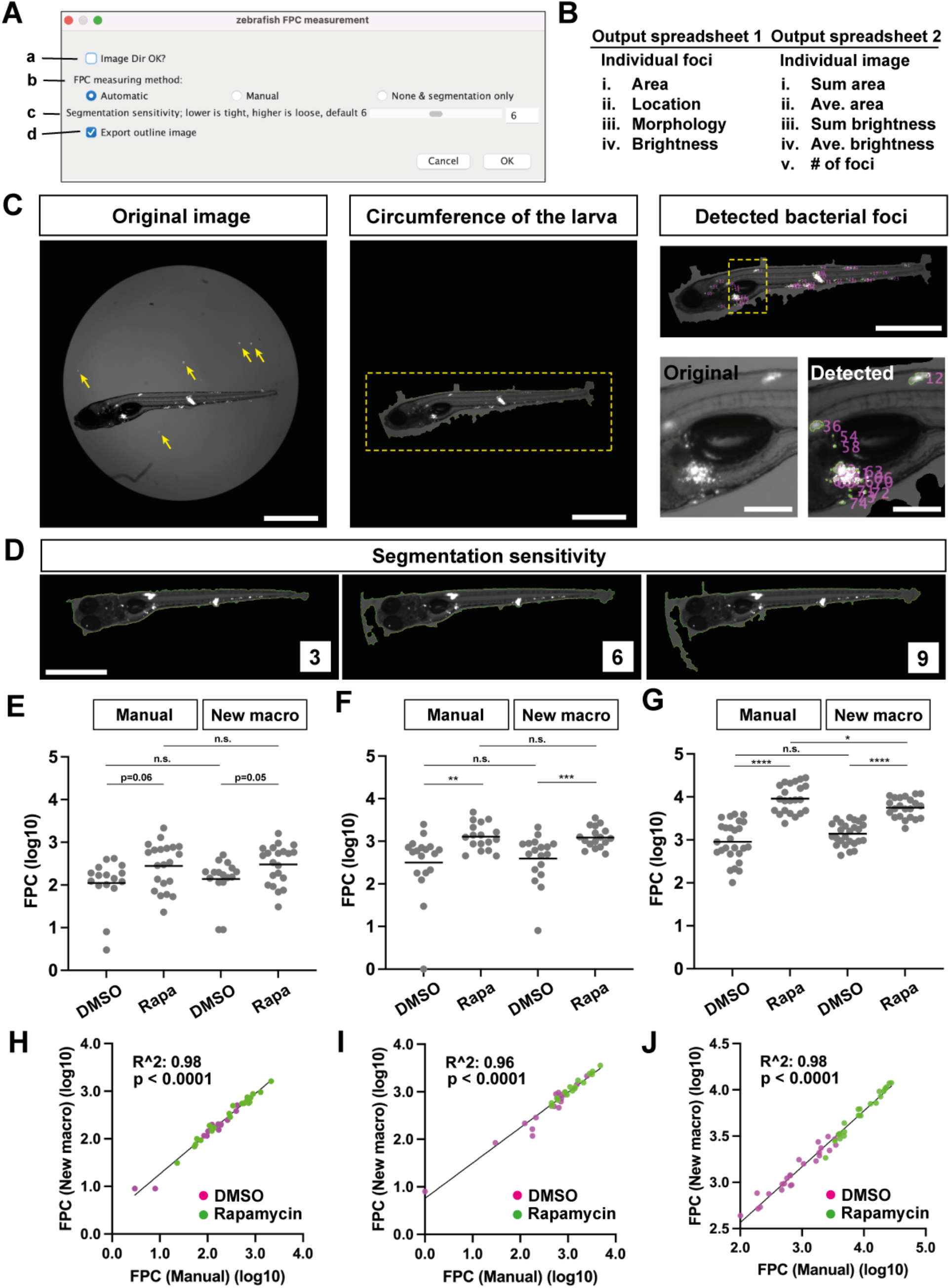
The newly developed image processing macro and its validation. (**A**) GUI of the macro. **a**, Selection of the folder containing the images to be processed. **b**, Selection of the image processing method to be used. We incorporated the conventional manual threshold determination (6) in the macro. **c**, Selection of Segmentation sensitivity. Default value is 6. See an example in Fig. 1D. **d**, Selection for either exporting segmented images or not. (**B**) Summary of the output files and parameters analyzed. See also **Supplemental Material 2**. (**C**) Example of the processed image and outcomes. **Left**: the original image (contrast adjusted). Yellow arrows: bright bacterial signals outside the larva. **Middle**: the result of automated detection of the circumference of the larva. A yellow dotted square is magnified in Right. **Right**: (top) the result of automated detection of the bacterial foci within the larva. (Bottom, left) the magnified original image taken from the location indicated with a yellow dotted square above. (Bottom, right) the magnified output image taken from the location indicated with a yellow dotted square above. The larva was 7 days post fertilization (dpf)/ 5 days post infection (dpi) injected with about 200 tdTomato-labeled Mm. Scale (left, middle, right top): 1 mm. Scale (right bottom): 200 μm. (**D**) Comparison of results using different Segmentation sensitivity values. The larva was 7 dpf/ 5 dpi injected with about 200 tdTomato-labeled Mm. Scale: 1 mm. (**E, F, G**) Comparison of FPC results obtained using the manual method and derived from the new macro. The larvae were 6 dpf/ 4 dpi (E, F) or 7 dpf/ 5 dpi (G) injected with about 20 (E) or 100 (F, G) tdTomato-labeled Mm, treated with 0.5% DMSO or 400 nM of Rapamycin in 0.5% DMSO from after the injection to 4 dpi (E, F) or 5 dpi (G). N = 16 and 21 (E), 18 and 17 (F), 26 and 22 (G) for the DMSO control and the Rapamycin treated groups, respectively. DMSO: 0.5% DMSO treated larvae, Rapa: 400 nM Rapamycin in 0.5% DMSO treated larvae. Each dot represents a larva. Mean values are indicated as horizontal lines. ***: p< 0.001, **: p< 0.01 and n.s.: p > 0.05 with Mann-Whitney test in (E) and (F). ****: p < 0.001, *: p < 0.05 and n.s.: p > 0.05 with two-tailed unpaired t-test (G). (**H, I, J**) The correlation of values derived from the two methods. (H), (I) and (J) were derived from (E), (F) and (G), respectively. R^2^ and p-value were analyzed with the simple linear regression and F-test, respectively.

To validate the new macro, we compared its outputs with those using our manual threshold-based FPC analysis, using the same image dataset. We selected datasets comprising Mm-infected zebrafish larvae, treated or not with the mTORC inhibitor rapamycin, which increases bacterial burdens (10) (detailed Materials and Methods are in **Supplemental Material 4**). We datasets where the manual FPC differences spanned ∼ one log for untreated larvae with the corresponding treated FPC’s being ∼ half to one log higher (Fig. 1E, F and G). The outputs from the manual method and the macro were highly correlated (Fig. 1H, I and J). They were also similar except in the last case, the manual FPC was a little higher in the treated group (Fig. 1G). Re-examination of the original images revealed that manual thresholding but not the automated macro had included bacteria with low intensity fluorescence (**Supplemental Material 5**, Supplemental Figure 2). Nevertheless, the automated macro identified the significantly higher bacteria burdens in the treated animals (Fig. 1G). Because manual FPC requires fluorescence thresholding based on identifying the lowest signal intensity that eliminates background pixels, it is subject to interoperator and interexperiment variability as illustrated in **Supplemental Material 5**. The automated macro has the benefit of removing subjectivity.

This automated, high throughout macro should be applicable for all fluorescent microbes studied in zebrafish larvae (7, 8, 11-14). The codes and instructions for the macro are freely available on GitHub (https://github.com/JaneliaSciComp/Zebrafish_96well_segmentation_mesure_FPC).

## Supporting information

Supplemental Material 1-5 combined

## Acknowledgements

We thank the Janelia Collaboratorium team, especially Misha Ahrens and Anoj Ilanges, and Ron Vale for encouragement and advice; N. Goodwin, R. Foster, and the University of Cambridge aquatic facility staff for zebrafish care; the Ramakrishnan lab for feedback on the macro. Funding: This study was supported by the HHMI Janelia Visiting Scientist Program, a Wellcome Trust Principal Research Fellowship (223103/Z/21/Z) and NIH MERIT award R37 AI054503 (L.R.); N.Y. was supported from EMBO postdoctoral fellowship (ALTF1130-2021) and JSPS Overseas Research Fellowships (202460397); M.E.B was supported by a Blavatnik Cambridge Fellowship, a Rothschild Yad Hanadiv Fellowship, Human Frontier Science Program (LT0029/2022-L) and the Israel National Postdoctoral Award for Advancing Women in Science.

## Author contributions

N.Y. H.O and L.R. conceived and designed the project. H.O and N.Y. conceived the macro. H.O. wrote the macro. N.Y. and M.E.B initially validated the macro with help from H.O. N.Y. and M.E.B. performed the experiments and analysis. N.Y wrote the paper with input from H.O. and M.E.B. L.R. edited the paper.

## Open access statement

For the purpose of open access, the author has applied a CC BY public copyright license to any Author Accepted Manuscript version arising from this submission. This work is licensed under a Creative Commons Attribution 4.0 International License.

## Competing interests

The authors declare no competing interest.

## Notes

### Competing Interest Statement

The authors have declared no competing interest.

