## Supplemental Material 1-5 combined for "An Image Processing Tool for Automated Quantification of Bacterial Burdens in Zebrafish Larvae"

### Manual method

The fluorescent bacteria shed from larval skin and other debris create bright spots outside the larval body (Supplemental Figure 1A1). They can interfere with the thresholding process during image analysis. Therefore, the manual method (6) requires manual blacking out of the space outside of the larva (Supplemental Figure 1A2). Also, this method requires a threshold value determined by each user (Supplemental Figure 1A3). The new macro in this study allows users to skip these processes (Supplemental Figure 1B).

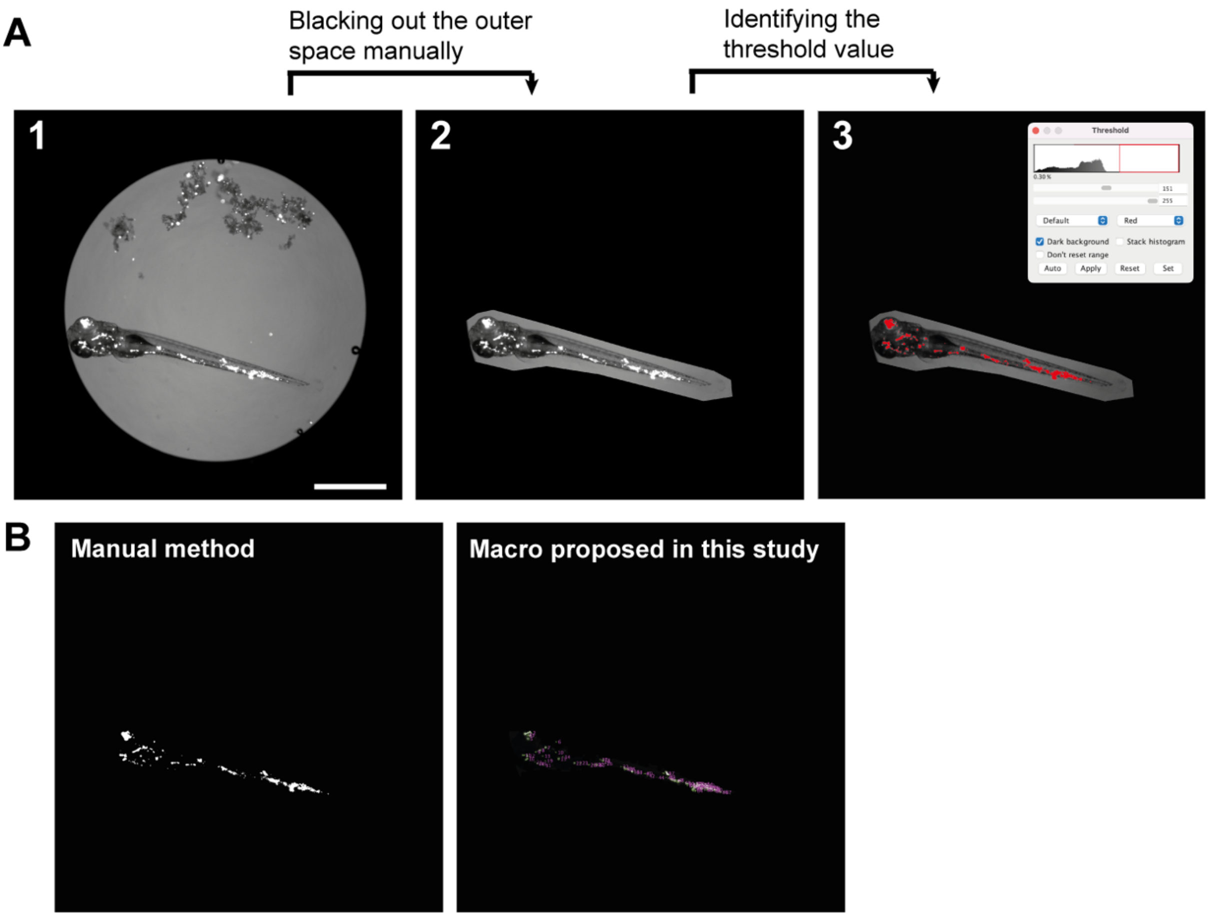


**Supplemental Figure 1: Explanation of the manual method and the output images from two methods.**

(**A**) The workflow of the manual image analysis. Scale: 1 mm. (**B**) Output images from the manual method (left) and from using the macro proposed in this study (right). The left image was created from A3. The right image was created using the macro with the input of A1.

**Supplemental Material 2**

**Output files**

When the macro is executed, it generates four types of output files.

The macro generates three output files for each individual image inside the folder containing processed images.

Those are:

**1**) A PNG image displaying the segmented outline of the larva (Fig. 1C, middle).

**2**) A PNG image showing segmented outlines and indexes of fluorescent bacterial foci (Fig. 1C, right).

**3**) A Microsoft excel spreadsheet containing measured parameters for segmented bacterial foci (Fig. 1B, left).

Finally, the macro generates **4**) A Microsoft excel spreadsheet as a summary of all processed images (Fig. 1B, right). This summary spreadsheet is created in the directory one level above the folder containing processed images. Conventional FPC values, identified as “Sum area”, can be easily found in this summary spreadsheet.

**Supplemental Material 3**

**Image analysis steps in the newly developed macro**

We opted for a series of relatively simple image processing techniques to accomplish this task. The typical input image, taken by a conventional widefield fluorescence microscope, utilizes both fluorescence and dim transmitted light to capture images of the larva and fluorescent bacteria within it simultaneously. We recommend using 16-bit images, instead of 8-bit images, to allow the macro to process with a wider range of signal intensities and to be able to make a segmentation even with the images with dim background.

To begin, the macro identifies and removes the circular border of the well from the 96-well image sing a rolling-ball-based background subtraction method. Subsequently, it locates the circumference of the larva through a multi-step process. Briefly, 1) the larval signal from the transmitted light is enhanced. 2) the threshold value for creating the larval binary mask is determined using a histogram-based standard deviation value. 3) the mask is evaluated based on shape, with an aspect ratio (length of the major axis/ length of the minor axis) required to be more than 4. The process 2 and 3 iterates until the process 3 is satisfied. If the algorithm fails to detect any objects meet the criteria, the macro rejects the image and proceeds to the next. Examples of such cases include images featuring an empty well or an unfocused larva. Although the outer border of the larva identified may not be perfect, it is normally sufficient to remove signals outside of the larva and proceed to the next step.

If the default Segmentation sensitivity (set at 6) does not produce a tight enough segmentation, users can opt for a lower value, as illustrated in Fig. 1D. In process 2 above, the macro searches for the threshold value to segment the larval object using the histogram of the gray values. It scans the histogram in increments of 10 (±5) gray values and calculates the standard deviation within each domain. The Segmentation sensitivity number determines the maximum standard deviation that the scanning process will allow before stopping and identifying the threshold gray value. For example, setting Segmentation sensitivity to 6 finds the domain where the maximum standard deviation is less than 6 and uses the maximum gray value in that domain for thresholding. Typically, the background has higher gray values, while the larval body has lower gray values, so selecting a lower Segmentation sensitivity usually results in a tighter segmentation (closer to the larval body). However, we also incorporated an aspect ratio-based decision, allowing the macro to iterate between these two methods. In rare cases, this may result in a tighter segmentation using a larger Segmentation sensitivity. Depending on the balance between the background and the larval body brightness, users may experiment various Segmentation sensitivity values to find the optimal setting for their purposes.

Once the larval object is determined, the macro detects fluorescence bacterial foci within it and measures their output parameters. This process also involves multiple steps. Within the larval mask, the minimum threshold value for the bacterial fluorescent foci is determined by the histogram-based standard deviation value. Fluorescent bacterial foci are identified through thresholding based on the local contrast, using the minimum thresholding value identified previously. Finally, the foci are indexed and their parameters are measured separately, with the values saved in the spreadsheets.

**Supplemental Material 4**

**Materials and Methods**

Ethics:

Zebrafish husbandry and experiments were conducted in compliance with a guideline from the UK Home Office using protocols approved by the Animal Welfare and Ethical Review Body of University of Cambridge.

Zebrafish husbandry, infections and treatment of Rapamycin:

Zebrafish embryos and larvae of the wild-type AB strain (Zebrafish International Resource Center: ZIRC) were used in experiments. All experiments were conducted in larvae such that their sex was undetermined. The general conditions for maintaining adult zebrafish were described before (10). Zebrafish embryos were collected and kept in egg water (0.18 g/L Instant Ocean Salt and 0.25 μg/mL methylene blue) at 28.5 °C without the day-night cycle. On 1 dpf, embryos to be used in experiments were transferred to 0.5x E2 medium (0.35 mM $NaHCO_3$, 0.5 mM $CaCl_2$, 0.025 mM $Na_2HPO_4$, 0.075 mM $KH_2PO_4$, 0.5 mM $MgSO_4$, 0.25 mM KCl and 7.5 mM NaCl) supplemented with 0.003% 1-phenyl-2-thiourea (PTU, Sigma).

For infections, 2 dpf larvae were dechorionated with 0.5 mg/mL of Pronase (Sigma). They were then anesthetized with medium containing 0.025% MS-222 (Sigma). Larvae were injected via the caudal vein using single-cell suspensions of Mm of known titer. Normally, the injection volume was 2-4 nL and the phenol red sodium salt (< 1% w/v prepared in PBS, Sigma) was used as a tracer for injection. Inoculums for each experiment were determined by injecting the bacterial sample onto the Middlebrook 7H10 plate supplemented with albumin, oleic acid, dextrose, Tween-80 and hygromycin B and later counting the number of colonies.

Rapamycin (Sigma-Aldrich, Cat# R0395; CAS: 53123-88-9) was dissolved in DMSO (Dimethyl sulfoxide, Fisher Scientific, Cat# BP231100; CAS: 67-68-5) as a 1 mM stock. Right after infection at 2dpf, 400 nM of Rapamycin with 0.5% DMSO or 0.5% DMSO was added directly in 0.5x E2 medium. We did not replace medium containing Rapamycin throughout the experiment.

Bacterial preparation:

*Mycobacterium marinum* M strain (ATCC #BAA-535) with expression of tdTomato under the control of the *msp12* promoter was used for all the infection experiments. These bacteria were grown at 33 °C in Middlebrook 7H9 medium (BD Difco) supplemented with albumin, Tween-80, oleic acid, and dextrose (Sigma-Aldrich). Hygromycin B (Cambridge Bioscience) was supplemented in culture to select the transgene positive Mm. The preparation of Mm single-cell suspensions was described before (6).

FPC measurements:

Widefield fluorescence larval imaging for performing FPC measurements was performed with a Nikon Eclipse Ti-E inverted microscope equipped with a 4x objective (NA 0.13) as previously described (10). Briefly, infected larvae were anesthetized with 0.025% MS-222 and subsequently ice-cooled for about 30 min to shrink their swim bladders. Larvae were then transferred to a 96 well half area cell culture microplate (Greiner Bio-One, Cat#675090) to take images. 16-bit fluorescence and bright field images were taken using a LED lamp with a 550 nm filter together with transmit light. Subsequently, all the post processing of images was performed using Image J/Fiji (9). The conventional FPC measurement was performed as follows. First, 16-bit images were converted into 8-bit images. Next, the outside of larva was manually blacked out for all the images. The threshold value, which was the lowest signal intensity that eliminated all the background signal, for the bacterial fluorescence was determined manually (100 for images analyzed in Fig. 1E and F and 80 for images analyzed in Fig. 1G). Finally, the previously developed fluorescence pixel counts macro (6) was ran with the determined threshold value in Image J/Fiji. The FPC measurements using the new macro was performed as follows. 16-bit original images were directly subjected to the macro with the default Segmentation sensitivity which was set at 6.

Statistical analysis:

Statistical analysis and visualization were performed on Graphpad Prism. Each statistical test used, mean and SD, the number of samples per group are indicated in the figure legend or in the panels. The Kolmogorov-Smirnov test and the F-test were performed to test for normality and for equality of two variances. Subsequently, the Mann-Whitney test and the unpaired two-tailed t-test were used to determine statistical significance between the mean values of two groups in Fig.1E and F, and in Fig. 1G, respectively. The simple linear regression was performed and the $R^2$ value and p-value, which was derived from F-test, was calculated.

**Supplemental Material 5**

**Comparing the new macro with the manual method**

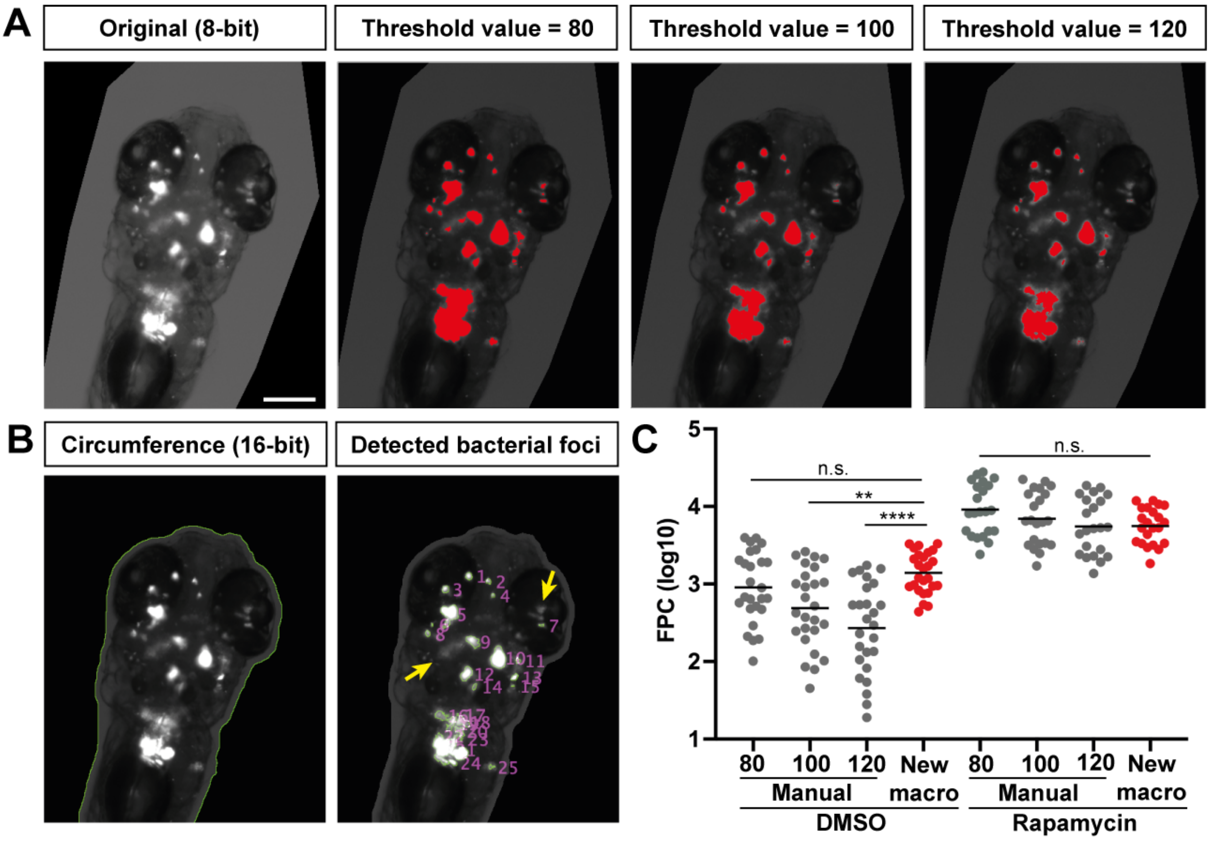


**Supplemental Figure 2: Comparison of the new macro with the manual method with various threshold values.**

(**A**) Detection of the bacterial foci with the manual method with various threshold values. The 7 days post fertilization/ 5 days post infection larva initially infected with about 100 tdTomato-expressing *M.marinum* (white) treated with 400 nM of Rapamycin for 5 days. Red areas are above the threshold values. Scale: 200 µm. (**B**) Detection of bacterial foci with the new macro. The new macro was used to analyze the same image as (A). Yellow arrows: low contrast bacterial signals that were not detected by the macro. (**C**) FPC values obtained with the manual method with the various threshold values and the new macro. The dataset for Figure 1E was reanalyzed. Horizontal lines: mean. n.s.: p>0.05, **: p<0.01, ****: p<0.0001 with Ordinary one-way ANOVA followed by Tukey's multiple comparison test.
